# Control of Quiescence and Activation of Human Muscle Stem Cells by Cytokines

**DOI:** 10.1101/2025.06.21.660874

**Authors:** Katharine Striedinger, Emilie Barruet, Elena Atamaniauc, Karla Linquist, Chris Knott, Andrew Brack, Jason H Pomerantz

## Abstract

Skeletal muscle homeostasis and repair depend on the activation of tissue resident stem cells called satellite cells. To understand the early molecular basis of human satellite cell activation, epigenomics, transcriptomics and protein analysis were performed in quiescent and activated human satellite cells. Cytokine signaling pathways were enriched in stimulated human satellite cells revealing a high cytokine enrichment, including CCL2, CCL20, CXCL8, IL-6, TNFRSF12A, ILR1, CSF-1 and FGF2. Functional roles of these observed changes are supported by *in vivo* experiments showing that chemokine inhibitors increase engraftment and regeneration capacity of human satellite cells xenotransplants. Cytokines, chemokines and associated signaling pathways in the early stages of human satellite cell activation may underlie disparate muscle responses in neuromuscular inflammatory and degenerative disorders and consequently are potential entry points for clinical applications towards muscle repair.

## Introduction

Skeletal muscle resident stem cells called satellite cells (MuSC), ensure muscle tissue homeostasis, repair and regeneration throughout life^1–3^. Quiescent and self-renewing satellite cells in mice and humans are characterized by the expression of the transcription factor PAX7, which is also expressed in activated MuSC but downregulated during activation and differentiation^4^. In homeostatic conditions, specific signals maintain MuSC quiescence. But upon muscle injury, MuSC transition from a G0 state to rapid activation and cell cycle re-entry (S phase), accompanied by orchestrated temporal, and spatial activation signals causing MuSC to express myogenic regulatory factors (MYF5, MYOD1, MYOGENIN) and progressive loss of PAX7 expression^1–5^. This leads to asymmetric divisions towards differentiation into muscle progenitor cells or myoblasts, migration, and fusion into multinucleated myofibers to regenerate muscle or symmetric self-renewal divisions for niche maintenance^6,7^.

While recent experiments in animal models have provided information about signaling mechanisms in the muscle niche ^4,6,8^, much of the environmental cuing that governs early MuSC activation remains less well understood, especially in humans. Furthermore, study of the impairment of satellite cell function across all categories of neuromuscular disorders found that 45% of myopathogenes (genes altered in neuromuscular disorders) are differentially expressed during initial hours of satellite cell activation^9^. A majority of these myopathogenes directly respond to PAX7 and MYOG.^9^

We recognized the importance of studying gene expression changes associated with the early activation process of human muscle satellite cells (Hu-MuSC). This required development of an experimental model of Hu-MuSC activation allowing for high throughput epigenomic characterization of activated compared to quiescent Hu-MuSC. By modeling Hu-MuSC activation in a reproducible experimental context, we found that in a short time window, Hu-MuSC undergo considerable epigenetic and transcriptomic changes. Among the most prominent groups of genes to change during early Hu-MuSC activation is cytokine signaling. Cytokines in satellite cells regulate quiescence and self-renewal, and chemokines (chemotactic cytokines) are involved in cell migration and proliferation in response to muscle damage.^10^

Here, we describe cytokine alterations upon Hu-MuSC activation and present novel observations of specific chemokine activity in activated Hu-MuSC along with experimental evidence of functional importance in muscle regeneration. Our findings also validate in human satellite cells several observations of satellite cell activation previously made in animal models and support functional impact of cytokines on satellite cells during activation. These findings have implications for understanding how satellite cell function is altered in human neuromuscular inflammatory and degenerative disorders.

## Results

### Experimental model of Hu-MuSC activation and associated epigenomic changes

Acknowledging that various muscle stimuli and injuries may differ in the exact satellite cell activation response, we sought to develop an experimental model of Hu-MuSC activation that would reasonably mimic common natural activation characteristics and would be reproducible. Such a model would be expected to induce some or all the canonical changes previously described to occur with satellite cell activation and would be reproducible. This model focuses on capturing events before cellular division and proliferation, focusing on the earlier stages of Hu-MuSC activation.

We used a combination of mechanical injury, ambient temperature and FGF2 enriched growth media to activate Hu-MuSC. Fresh muscle biopsies were macro-dissociated and incubated at 37 C to activate the satellite cells (detailed in Methods and in previous publications from our laboratory^11,12,13^). The remaining muscle fraction from the same donor muscle sample, the control sample, was kept at 4 C and not macro-dissociated until just before Hu-MuSC isolation (**Figure 1a**). Hu-MuSC isolation was done for control and activated samples at the same time, using standard human satellite cell markers (CD29, CD56 and CxCR4) and sorted by FACS (**Figure 1a**).

**Figure 1.**
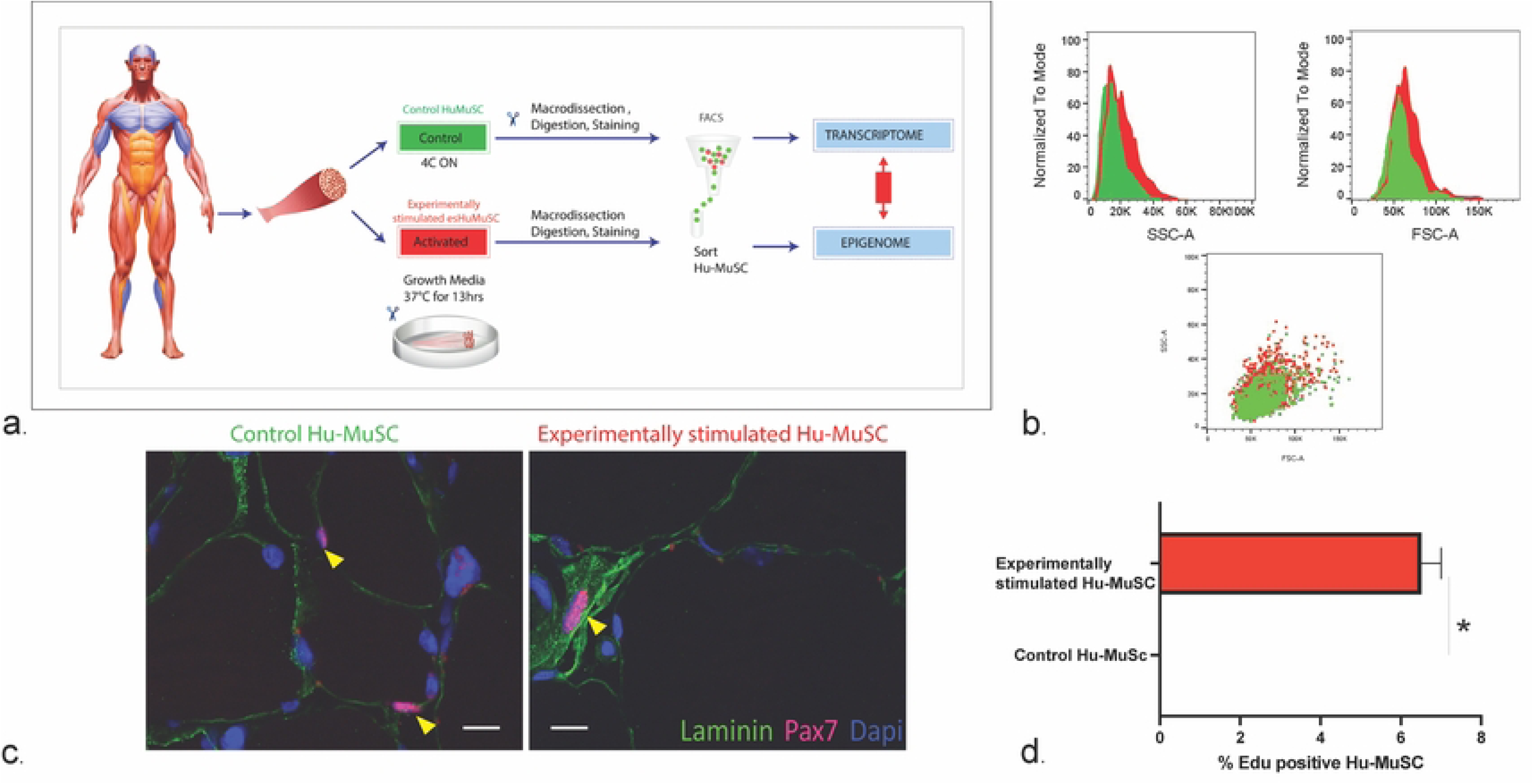
Model of human satellite cell activation. **(a)** After collection, human muscle biopsies were either kept at 4°C overnight until dissociation (Control) or immediately dissociated and incubated with growth media at 37°C overnight until further mechanical and enzymatic dissociation (Experimentally stimulated). Then, human muscle satellite cells (Hu-MuSC) were sorted for CXCR4+/CD29+/CD56+ using fluorescence activated cell sorting (FACS) and used in downstream experiments, for example single cell transcriptomics and epigenomics. **(b)** An increased Side (SSC-A) and Forward (FSC-A) scatter in the stimulated Hu-MuSC (red) compared to of control Hu-MuSC (green) reveal an increased cell size and granularity once the Hu-MuSC undergo activation. **(c)** Hu-MuSC from *in situ* activated human muscle show an increase in size (yellow arrowhead) identified with PAX7 (red) immunostaining compared to control muscle Hu-MuSC. Scale bar: 10 uM. **(d)** Higher percentage of EdU labeled Hu-MuSC in the activated muscle (6.5 ± 0.2%) compared to non-labeled not dividing quiescent control Hu-MuSC 24 hrs after FACS sorting.

Acknowledging possible differences between Hu-MuSC activation in experimental models and natural settings, we term activated Hu-MuSC, experimentally stimulated Hu-MuSC (esHu-MuSC). esHu-MuSC were increased in size as shown in the forward and side scatter images compared to controls (**Figure 1b**). High magnification images of the control and activated myofiber fragments also show increased size of the PAX7 immunolabeled esHu-MuSC (**Figure 1c**). Only esHu-MuSC, were positive for Edu (6.5 ± 0.2%) within the first 8 hrs after sorting compared to controls (**Figure 1d**). Stimulated Hu-MuSC and control Hu-MuSC express different levels of PAX7 and MYOD1 expression after FACS isolation at the epigenomic (**Figure 2f**), transcriptomic (**Figure 3a,b**) and protein levels (**Figure 4a**) confirming these are satellite cells in early stages of activation. Thus, esHu-MuSC express expected characteristics of activated MuSC supporting this model of satellite cell activation.

**Figure 2.**
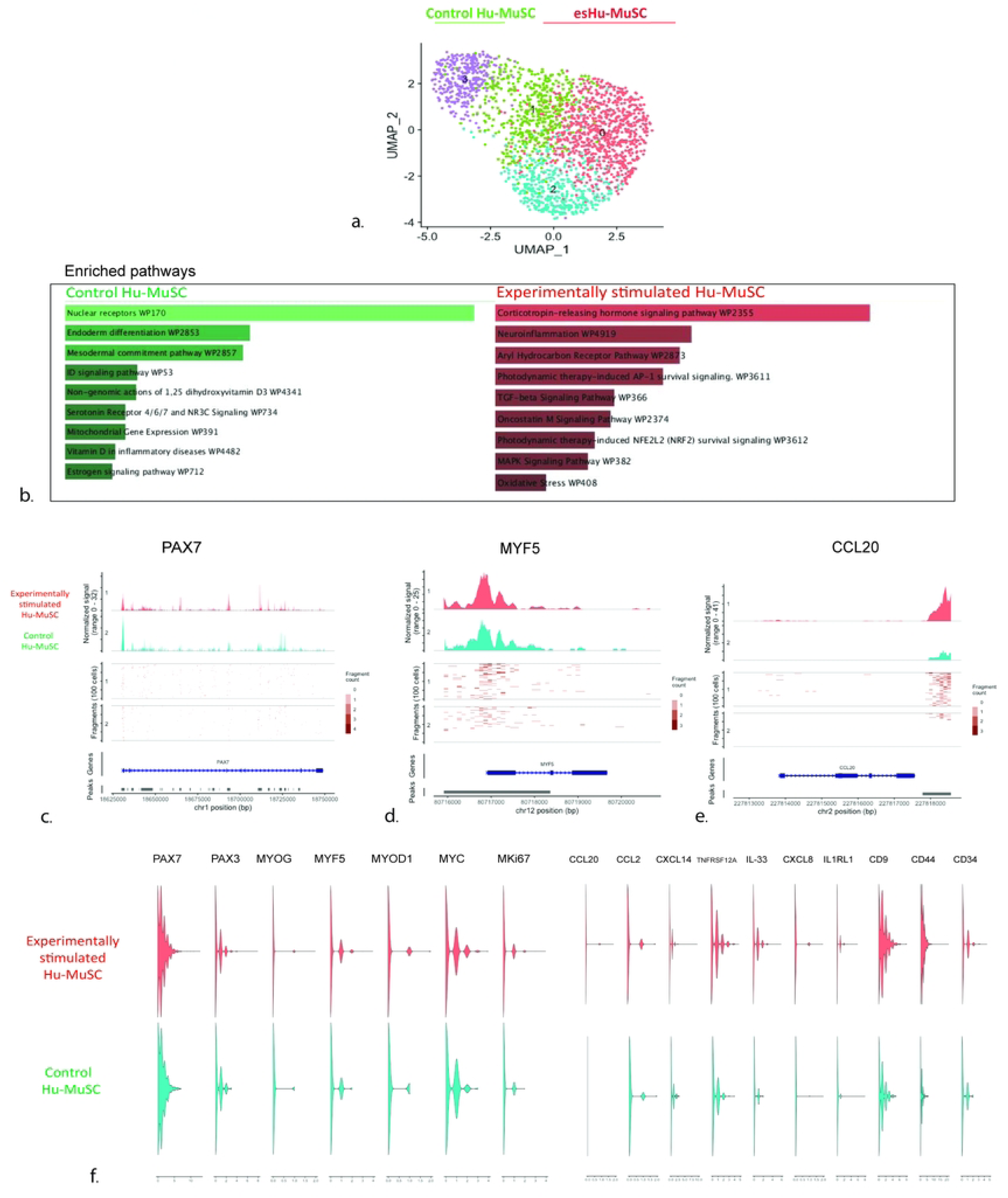
Single-nuclei chromatin accessibility assay (snATAC) in control and experimentally stimulated human satellite cells from the vastus muscle of a 56-year-old man**. (a)** UMAP representations of snATAC-seq of control and esHu-MuSC. Individual dots represent single cells, here in total 9,232 Hu-MuSC nuclei, distributed in 7,850 from esHu-MuSC, and 2,411 from control Hu-MuSC. A clear epigenetic shift distinguishing control from esHu-MuSC becomes apparent in the resulting four Hu-MuSC clusters (resolution of 0.8). **(b)** Top differentially expressed pathways based on the highest differential peak expression between control and stimulated Hu-MuSC sorted by p-value using WikiPathways^14^. Note the TGF-B, Oncostatin-M and MAPK, among the other listed pathways that are significantly enriched in the activated Hu-MuSC. **(c-f)** Violin plots showing the distribution of chromatin fragments using scATAC, on esHu-MuSC and control Hu-MuSC. Open chromatin violin plots showing presence of PAX7 (**c**) in both esHu-MuSC and controls while increased fragments of Myogenic factors like MYF5 (**d**), MYOG, MYC and MYD1(**f**) in the esHu-MuSC compared to controls. **(e,f)** Increased distribution and frequency of chemokine open chromatin fragments including CCL20 **(e)**, CCL2, IL-33 in esHu-MuSC compared to controls **(f).** These results show significant epigenetic cytokine peaks during activation, preceding overall changes in muscle stem cell transcription factors and myogenic regulatory genes in human satellite cells.

**Figure 3.**
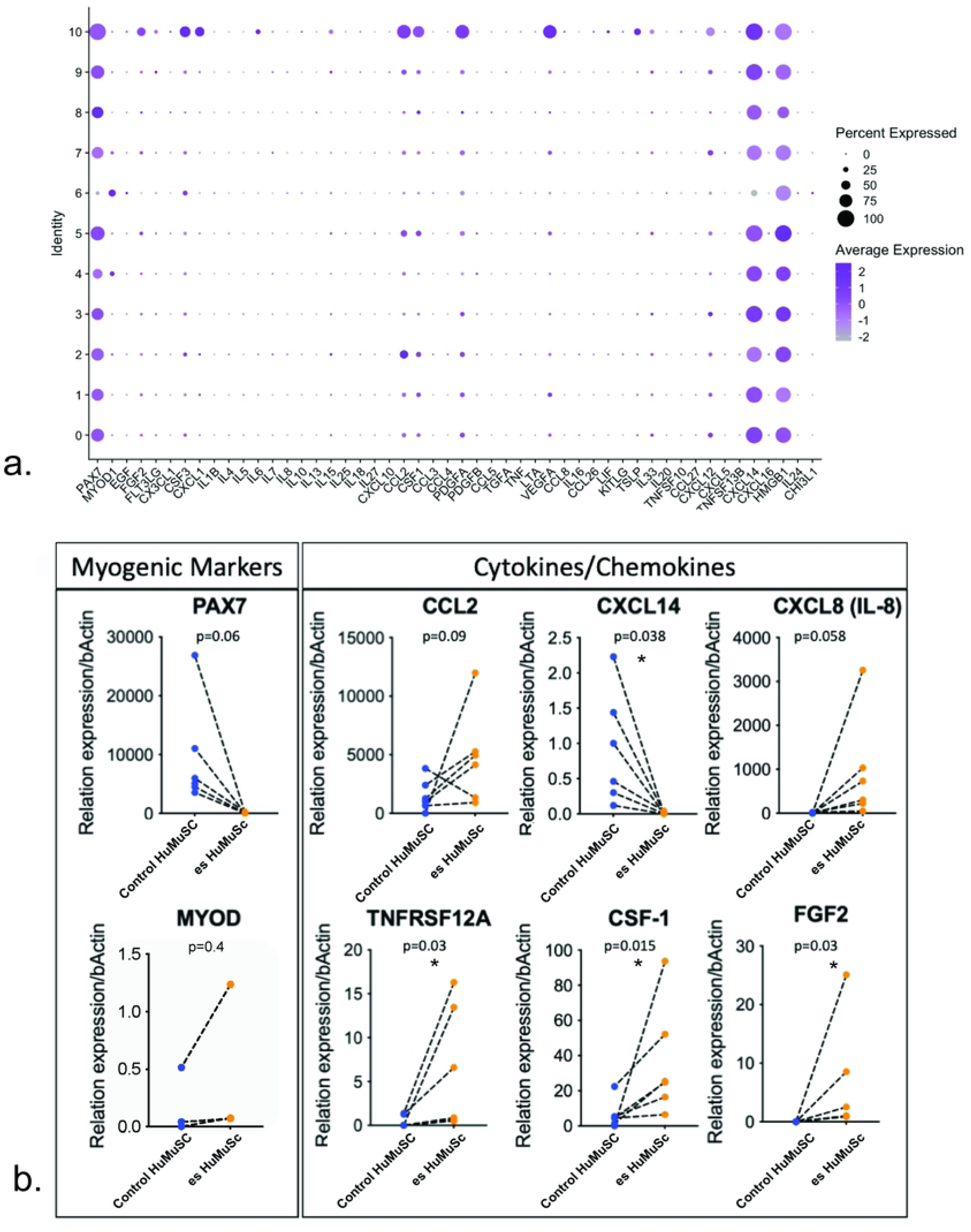
Differential temporal expression of cytokines and chemokines ligands and receptors during human satellite cell activation. **(a)** We evaluated cytokine and chemokine transcription of Hu-MuSC from twelve different donors, in three muscles: vasti, pectoralis and rectus muscle. As expected, the majority of Hu-MuSC in these samples express high levels of *PAX7* transcripts and lower levels of *MYOD1* Hu-MuSC. Cytokine transcripts such as, *CCL2, CXCL14* and *HMGB1* are abundantly expressed among all Hu-MuSC clusters from unstimulated muscle. Cluster 10 exhibits the highest percentage of Hu-MuSC expressing cytokines, chemokines transcripts and growth factors, including *FGF2, CSF3, CXCL1, CCL2, CSF1, PDGFA, VEGFA, CXCL12*. **(b)** To further evaluate cytokine expression during satellite cell activation we compared cytokine transcripts expression using RT-PCR in control and experimentally stimulated Hu-MuSC (esHu-MuSC). Consistent with our epigenomic findings, there is a tendency of *PAX7* transcripts to be rapidly downregulated after activation, and elevated levels of *MYOD1* upon activation. The tendency for chemokines like *CCL2 and CCL8* to be upregulated in the esHu-MuSC, as well as a significant upregulation of TNFRSF12A, CSF-1 and FGF2 in the esHu-MuSc compared to controls. In contrast CxCL14 was found significantly downregulated in esHu-MuSC compared to controls (Wilcoxon test *P*-values are *<0.05).

**Figure 4.**
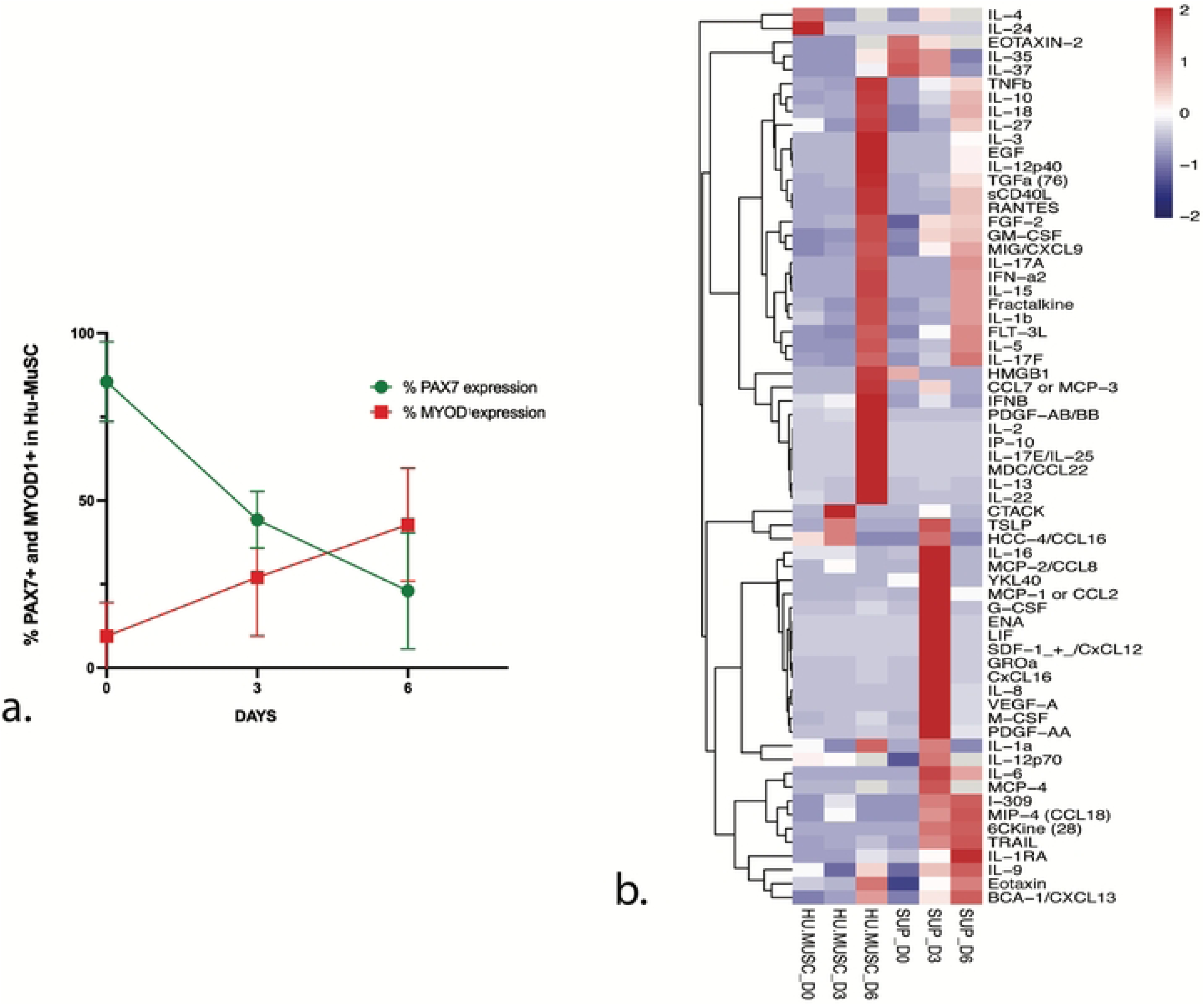
The expression of PAX7 and MYOD1 correlate with a differential expression of diverse chemokines in Hu-MuSC. **(a)** Quantification of protein expression for PAX7+ and MYOD1 positive Hu-MUSC at day 0, 3 and 6 in vitro, showing an inverse correlation as progressively PAX7 expression decreases and MYOD1 expression increases in Hu-MuSC as they activate *in vitro*. using linear regression PAX7 slope varies significantly p= 0.0001 and MYOD1 slope also varies significantly p=0.05 (**b**) Heat map displaying the expression levels of all cytokine’s protein detected in Hu-MuSC from collection as they activate progressively at day 0, day 3 and day 6 *in vitro*, using the high throughput proteomic Multiplex assay. Out of 93 screened we found 76 cytokines in the Supernatant of Hu-MuSC, with the highest concentrations including IL-8, CCL2, GROa and M-CSF. Notice the sequential expression of cytokines at different time points, with early expression at day 0 of IL-4 and IL-24 and in the IL-35, IL-37 and Eotaxin-2 in the supernatant at day 0. And later at day 3 the peak of expression in the supernatant of CCL2, CCL8, LIF, IL-8, IL-6 peaks of protein expression occur at day 3 *in vitro* and other cytokines being mostly expressed later at day 6 in both Hu-MuSC lysates and supernatant. Each pair of observations connected with a dotted line was derived from an independent human muscle sample.

To understand molecular epigenetic modifications that may control the observed cellular changes of early activation, we compared the open chromatin regions, the significant peaks, and the cellular pathways, differentially expressed in control Hu-MuSC and esHu-MuSC, using the single cell Assay for Transposase Accessible Chromatin (scATAC). Hu-MuSC were isolated from a 56-year-old male vastus lateralis, to which we applied the described activation protocol and collected stimulated and control Hu-MuSC (**Figure 2**). Nuclei were isolated and processed for library preparation and sequencing. Sample performance in single cell ATAC assay in control and esHu-MuSC using the DNA accessibility assay demonstrated QC metrics including: Nucleosome signal QC metric distribution, DNA fragment length distribution the strength of the nucleosome banding pattern, the insert size distribution and the transcriptional start site (TSS) enrichment, within the recommended values. Low-quality cells and duplets were removed based on these QC metrics.

The resulting dataset of 9,232 cells distributed in 7,850 from stimulated Hu-MuSC and 2,411 from the control Hu-MuSC. Counterintuitively, the median fragments per cell were higher in the control Hu-MuSC (14,631) compared to the stimulated Hu-MuSC (5,713). This observation suggests a high level of open chromatin in control non-stimulated Hu-MuSC, supporting the concept of an active regulation of the quiescent state.

After data normalization and linear dimensional reduction, we constructed a low-dimensional visualization of the DNA accessibility assay using uniform manifold approximation and projection (UMAP). The UMAP shows a clear shift in chromatin conformation upon Hu-MuSC stimulation (**Figure 2a**), distinguishing control clusters (mainly distributed in cluster 1 and cluster 3) from the esHu-MuSC, represented mostly in clusters 0 and 2 (**Figure 2a**).

We identified significantly differentially expressed peaks (p values < 0.05) between control and esHu-MuSC. We identified 47 significantly different genomic accessible peaks from which 10 corresponded to non-coding RNA. Remarkably, 8 of 37 coding regions, corresponded to known cytokines, chemokines, immune system mediators and growth factors. We then performed cellular pathway analysis based on the enriched peaks and open chromatin regions in control and esHu-MuSC. Genome wide pathway analysis revealed increased activity of interleukin and immune system pathways in esHu-MuSC. Top differentially expressed pathways based on the highest differential peak expression between control and esHu-MuSC were sorted by p-value using WikiPathways^14^ (**Figure 2b**). While the control Hu-MuSC enrichment is on nuclear differentiation and mitochondrial gene expression pathways, the esHu-MuSC exhibit enrichment of corticotropin releasing hormone, neuroinflammation, TGF-beta signaling, Oncostatin and MAPK signaling pathways (**Figure 2b**).

PAX7 open chromatin was present in both stimulated and control samples, but upon stimulation, PAX7 open chromatin first peaks decrease, especially at the transcription start site (TSS) region of the PAX7 gene (**Figure 2c**), while MYF5 transcription factor open chromatin peaks increase in the esHu-MuSC (**Figure 2d**).

Since many of the altered cellular pathways involve cytokines and their effectors, we decided to further investigate the most significant changes. The CCL20 (**Figure 2e**), monocyte chemoattractant protein-1 (MCP-1, CCL2) and the C-C motif chemokine ligand 8 (CCL8) also known as MCP-2, have higher peaks in esHu-MuSC compared to paired controls (**Figure 2e,f**). MKi67 has more open chromatin fragments in esHu-MuSC compared to controls (**Figure 2f**). esHu-MuSC ATAC data reveals that inflammatory cytokines including IL-33 and CXCL8 (**Figure 2f**), and the receptors TNFRSF12A (TWEAK) and IL1RL1, as well as the antigen receptors CD14, CD44, CD9 increase upon stimulation (**Figure 2f**). These findings demonstrate a broad engagement of the known cytokine, chemokine and immune modulating pathways immediately after Hu-MuSC stimulation.

### Cytokine transcriptome changes upon Hu-MuSC activation

The open chromatin results prompted us to confirm gene expression using analysis of single cell RNA from freshy isolated Hu-MuSC. We have previously shown that Hu-MuSC from healthy muscle biopsies exist in distinct transcriptional clusters that differ in their gene expression profiles^12^. Within a specific muscle sample, the majority of Hu-MuSC express patterns consistent with quiescence, but have distinct expression profiles of unclear significance, and some express genes associated with activation^12^.We evaluated cytokine and chemokine transcription of Hu-MuSC from 12 human muscle samples. As expected, the majority of Hu-MuSC in these samples express high levels of *PAX7* transcripts and lower levels of *MYOD1* Hu-MuSC (**Figure 3a**). Cytokine transcripts such as, *CCL2, CXCL14* and *HMGB1* are abundantly expressed among all Hu-MuSC clusters from human muscle (**Figure 3a**). Cluster 10 exhibits the highest percentage of Hu-MuSC expressing cytokines, chemokines transcripts and growth factors, including *FGF2, CSF3, CXCL1, CCL2, CSF1, PDGFA, VEGFA, CXCL12* (**Figure 3a**).

To further evaluate cytokine expression during satellite cell activation we used RT-PCR from paired control and experimentally stimulated Hu-MuSC. Each pair comes from a single muscle donor (**Figure 3b**). Consistent with our epigenomic findings, *PAX7* transcripts are rapidly downregulated after activation, giving rise to elevated levels of *MYOD1* upon activation (**Figure 3b**).

There is a tendency for chemokines like *CCL2* and *CCL8* to be upregulated in the esHu-MuSC, compared to controls and a statistically significant increase of *TNRSF12A*, *CSF-1* and *FGF2* in stimulated Hu-MuSC compared to controls (**Figure 3b**). These quantitative increased tendency of these cytokine transcripts in esHu-MuSC was also observed in the open chromatin data. Note *CxCL14* is significantly downregulated upon activation in esHu-MuSC(**Figure 3b**).

To further characterize the activation state of the cells in this assay, protein expression was evaluated by immunostaining. The percentage of PAX7+ and MYOD1+ Hu-MuSC *in vitro* over the first 6 days was quantified. PAX7 significantly decreases to less than half in the first 3 days while MYOD1 significantly and progressively increases over 6 days (**Figure 4a**), confirming that this assay captures the early temporal window of Hu-MuSC activation and progression towards differentiation.

To investigate the cytokine protein expression during culture activation, we used a high throughput proteomic multiplex assay to identify the cytokine ligands and receptors expressed in Hu-MuSC from three different muscle samples. With the Human Cytokine/Chemokine Array (Eve technologies), we tested 92 different cytokines in the supernatant and in the cellular lysates of Hu-MuSC collected at day 0, at day 3 and at day 6 after Hu-MuSC were plated under growth conditions. After background subtraction analysis, 35 cytokines were found to be expressed at the protein level in Hu-MuSC, with varying concentrations during activation in culture (**Figure 4b**).

The cytokine protein concentration in cell lysates and supernatant is represented in a Heat map (**Figure 4b**) displaying the expression levels at different times points. The earliest expressed cytokines were IL-4 and IL-24, detected in lysates at day 0 while Eotaxin-2, IL-35, IL-37 were the earliest detected cytokines in the supernatant (**Figure 4b**). The expression of IL-8, YKl40, CCL2, GROa and M-CSF is apparent on day 3 in supernatant. Low levels of ten more cytokines were also detected in the cellular lysates, including IL-4, IL-24 and CCL16 as early as day 0 (**Figure 4b**). Noticeable, CCL2, IL-8 and IL-6 show highest expression in Supernatant at day 3.

We observed the maximum expression of TGF-B, CCL7, HMGB1 in Hu-MuSC lysates at day 6. A singular cytokine, HMGB1, was found to be expressed in Hu-MuSC at higher concentrations than all the other cytokines (**Figure 4b**). The expression of PAX7 and MYOD1 correlate with a differential expression of diverse cytokines and chemokines in Hu-MuSC (**Figure 4a,b**)

### Chemokine gain and loss of function modulates engraftment after Human Satellite Cell Xenotransplantation

Our expression data suggest that chemokines may play a functional role during Hu-MuSC activation. Moreover, satellite cell activation and muscle repair involve migration from the niche at the basement membrane to the center of the myofiber to initiate myofiber regeneration. This migration could be expected to be controlled by chemokine activity. We identified CCL2 as one of the most abundantly present chemokine transcripts in Hu-MuSC and found that it is actively upregulated during early activation. Thus, CCL2 was selected as a candidate target to explore the effect of gain and loss of function on engraftment and myofiber formation by Hu-MuSC *in vivo*.

We treated immunodeficient mice with a daily dose of either, CCL2 agonist or Bindarit or vehicle controls for 7 days, after Hu-MuSC transplantation in the Tibialis Anterior (TA) (**Figure 5a**). We used the CCL2 antagonist Bindarit, an indazolic derivative, to inhibit CCL2^15^. The effect of Bindarit is thought to be mediated by the downregulation of the NF-κB pathway^15,16^ and TGF-B signaling, inhibiting CCL2, and also affecting CCL7 and CCL8 ^15^.

**Figure 5.**
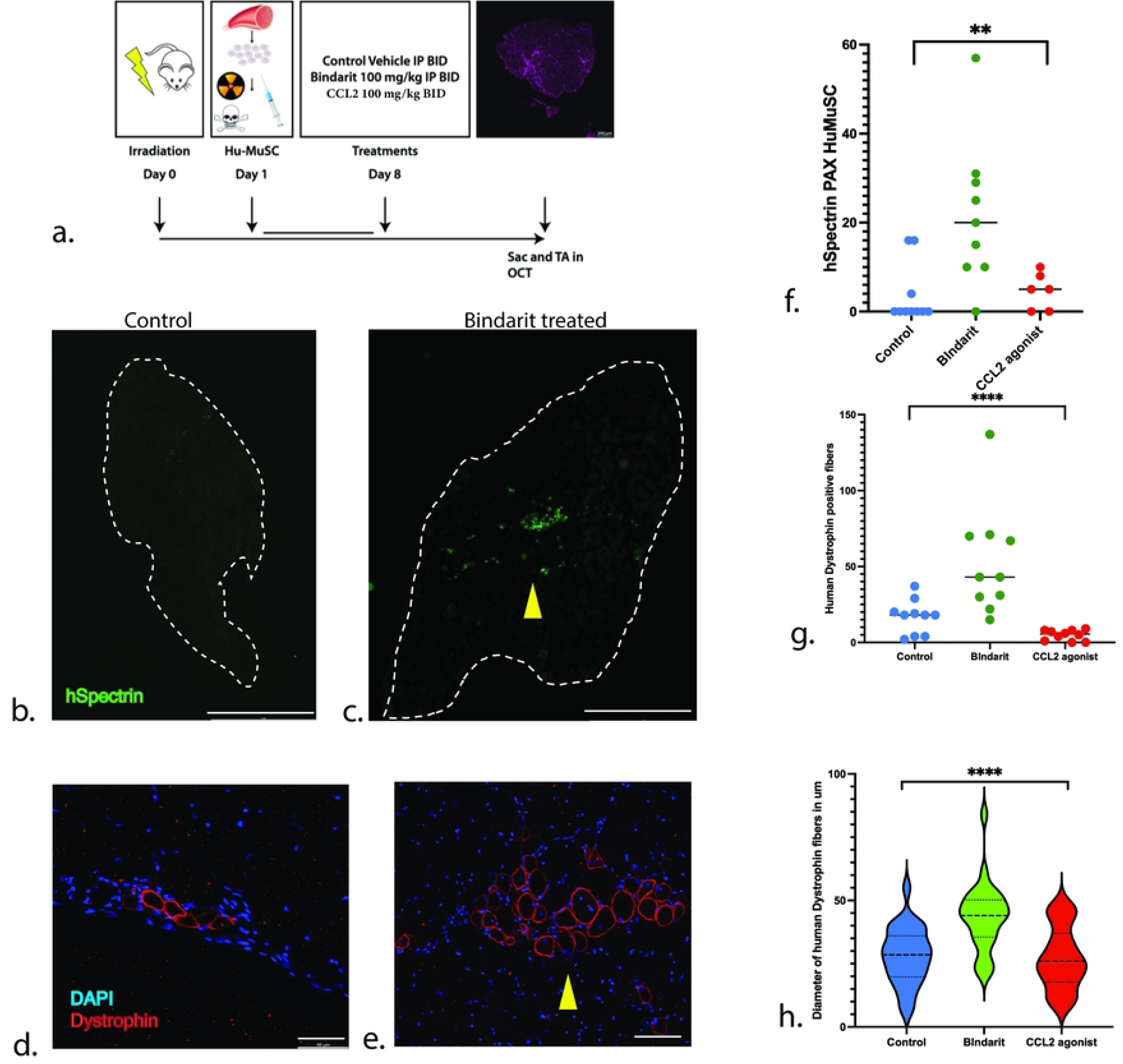
Xenotrasplants of Hu-MuSC in TA of mice. **(a)** Model for xenotransplants of Hu-MuSC in the TA of NSG mice, treated with the cytokine inhibitor Bindarit, or a CCL2 agonist, or vehicle control. (**b-e**) Sections of TA showing a representative example of Control (**b,d**), Bindarit (**c,e**) treated mice. By means of immunostaining, we observe a high number of human spectrin **(b,c,f)** and Dystrophin positive fibers **(d,e,g)** in the Bindarit treated muscle (yellow arrowheads) compared to controls (**g**). The diameter of the newly formed human fibers is significantly higher in the Bindarit treated compared to controls (**h**). N=10 mice for each group. Scale bar represents (**b, c**: 1 mm); (**d, e**: 50 μm) Statistical analysis was performed using paired Student’s *t-*tests; *P*-values are *<0.05, ***<0.001.

According to our single cell chromatin data, these chemokines are all significantly enriched in stimulated human satellite cells. Three weeks after transplantation of 2,000 Hu-MuSC, the TAs were collected and the number and diameter of the human Dystrophin positive fibers as well as the number of xenotransplanted Hu-MuSC expressing PAX7+ were quantified: a significantly higher number of human Spectrin (**Figure 5b,c,f)** and Dystrophin positive fibers (**Figure 5d,e)** were observed in the Bindarit treated muscle compared to controls (**Figure 5g**). The diameter of the newly formed human fibers is also significantly higher in the Bindarit treated group compared to controls (**Figure 5h**). These results show a more robust cluster and higher number of human fibers in the Bindarit treated group compared to controls and CCL2 agonist (**Figure 5b-h**) suggesting that Bindarit promotes differentiation and associated engraftment of Hu-MuSC.

Co-Immunostaining of PAX7 and human Spectrin were used to identify undifferentiated Hu-MuSC. While in controls, Hu-MuSC are sparse as in prior studies in quiescent muscle, in the Bindarit treated group there were more frequent Hu-MuSC PAX7+ and hSpectrin+ cells, suggesting that Hu-MuSC survive, returned to quiescence and/or self-renew more readily in a chemokine-depleted environment (**Figure 5f**). These results support the idea that reducing the levels of chemokines in the host muscular niche by means of exogenous agents, can result in higher engraftment after transplantation and greater myofiber formation capacity (**Figure 5a-h**).

## Discussion

In this study, we examined the epigenetic events of the transition of human satellite cells from quiescent stem cells, into activated proliferating muscle-stem cell progenitors, a process essential for proper muscle maintenance and regeneration. Our findings identify a complex network of cytokines, chemokines, ligands, and associated pathways acting simultaneously during the early activation of human satellite cells.

Our experimental model of early Hu-MuSC activation recapitulates previously described changes of activation, including increase in size, granularity and EdU uptake ^4,17^. It has been described that the low metabolism of quiescent MuSCs is primarily dependent on mitochondrial fatty acid oxidation and oxidative phosphorylation to promote epigenetic modifications that repress myogenic transcription programs^4^ and conserve the stemness of MuSC. In contrast, activated MuSC display a different metabolic phenotype as they increase in cell size and shift toward anaerobic glycolysis^4^. This supports a metabolic environment that allows for rapid biosynthesis, therefore, supporting growth and proliferation.

Despite the metabolic shift during activation, our single cell open chromatin data showed that the median fragments per cell were almost three times higher in the control Hu-MuSC compared to the experimentally stimulated Hu-MuSC, in support of the concept that quiescence is not a passive process but a highly transcriptionally active cellular state under tight regulation to decrease cellular stress and ensure long term maintenance^17–19^. This is consistent with chromatin data studies in MuSC^20^, where the chromatin of quiescent MuSC is largely permissive and becomes more restricted due to increased H3K27 upon MuSC activation, yielding a landscape with many genes enriched in quiescent MuSC but rapidly downregulated upon activation^20^.

MuSC quiescence is associated with PAX7 expression and inhibition of MYF5 and MYOD1, which in turn are recognized hallmarks of MuSC activation and myogenic commitment^4^. The early tissue activation model for satellite cells used here allowed observation of some of these earliest changes. Accordingly, we confirmed the expected observation that human unstimulated satellite cells have higher *PAX7* open chromatin fragments than stimulated Hu-MuSC but stimulated Hu-MuSC also contain abundant *PAX7* fragments while expressing higher levels of *MYF5, MYOD1* and *MYOG* than controls*. MYF5* is one of the first myogenic regulatory factors to increase with early satellite cell activation^21^. These epigenetic events were detected at the single cell level in this experimental system before detectable changes in other myogenic regulatory factors such as *MYC* and later on *MHC* take place, which characterize states further downstream in myogenic commitment, differentiation and myofiber fusion^4,21^.

Studies in mice and humans have identified at least two major subgroups of satellite cells by single cell RNA-seq^4,12,22^. MuSC close to quiescence express high levels of *PAX7*, *Hes1*, *Col2A*1 and genes related to cell cycle arrest and stress resistance, while MuSC in early activation express *MYOD1*, decreased *PAX7* expression and enrichment of genes involved with protein synthesis^22^. Although almost 100% of Hu-MuSC express PAX7 after isolation from adult human skeletal muscle^13^, they rapidly lose PAX7 expression over next 3 days *in vitro*, while increasing expression of MYOD1, mimicking the activation of satellite cells towards myoblasts^13^. This marks a short window of quiescence for Hu-MuSC *ex vivo* in which we identified extensive changes in cytokine and chemokines transcripts and protein expression. The increase in open chromatin configuration of cytokines in our scATAC experiments correlate with most of the changes in the myogenic regulatory factors during activation of Hu-MuSC. Therefore, cytokines are one of the earliest groups of genes to change in the activation of Hu-MuSC and likely do not depend on the myogenic regulatory factor cascade to do so. These findings imply that important regulators and effectors of MuSC behavior act in parallel and possibly independent of canonical myogenic pathways during activation and regeneration.

As Hu-MuSC become activated, the chromatin regions of cytokine *TNFRSF12A* become highly exposed. TNFRSF12A (TWEAK receptor) and the TNF receptor superfamily member fibroblast growth factor-inducible 14 (Fn14), have emerged recently as a pivotal axis for shaping both physiological and pathological muscle responses to acute or chronic injury and disease and in regulating skeletal muscle mass and metabolism^21,23,24^. TWEAK expression results in enhanced proliferation of mouse muscle myoblasts^23^. TWEAK also inhibits satellite cell self-renewal by inhibiting Notch signaling and stimulates the mitogen-activated protein kinase (MAPK) and canonical and non-canonical nuclear factor-kappa B (NF-κB) signaling pathways in myoblasts^21^. FGF (fibroblast growth factors) and MAPK (mitogen-activated protein kinase) signaling pathways also regulate MuSC proliferation and myogenic commitment. FGF signaling from myofibers, satellite cells and fibroblasts controls differentiation, whereas p38 MAPK has been implicated in both self-renewal and differentiation. Our human chromatin supports these findings in mice as a conserved pathway in human muscle, as we also observed TWEAK receptor, MAPK, NF-κB and cytokine signaling pathways enriched in stimulated human satellite cells.

Several transcription factors, cytokines and ligands previously found to have roles in muscle stem cell activation and repair were identified in our experiments in human cells. KLF10, an effector of TGF beta signaling has a functional role in proliferation and differentiation of a variety of tissues and loss of KLF10 in normal and mdx mouse skeletal muscle results in increased fibrosis and decreased grip strength^25^. IL-6 and KLF15 were among the predicted upstream regulators for later activation in a bulk chromatin accessibility study comparing human cultured myoblasts to satellite cells^26^. In animal studies, cytokines that have been described to play active roles in quiescence include the cytokine CXCL14, an endogenous inhibitor of myoblast differentiation and skeletal muscle regeneration^27^. The CXCL12–CXCR4 axis is known to maintain stem cell quiescence and CXCL14 depletion has been implicated in cell cycle withdrawal and acceleration of myogenesis^27^. Oncostatin M (OSM), a member of the interleukin-6 family of cytokines, is a potent inducer of murine muscle stem cell quiescence acting though the hippo pathway and yap to stimulate self-renewal^28^. In contrast, our human epigenetics data shows that Oncostatin M pathways are enriched in stimulated human satellite cells.

Several inflammatory myopathies have been associated with CCL2 (MCP-1), MCP-3 and RANTES ^29–31^, all these differentially expressed in the first 72 hrs in our Hu-MuSC in our experiments. CCL2 is one of the key chemokines implicated in the regulation of migration and infiltration of monocytes/macrophages. CXCL12 and CCL2 has been shown to be increased in idiopathic inflammatory myopathies^32^; here we described CxCL14 and CCL2 chemokines differentially expressed during Hu-MuSC activation.

Our *in vivo* xenograft experiments show that by decreasing CCL2, CCL7 and CCL8 activity with Bindarit, transplanted Hu-MuSC develop more and larger regenerated human myofibers. These data indicate that the engraftment of Hu-MuSC and myofiber formation is enhanced by transient inhibition of inflammatory cytokines. It is possible that lowering the expression of inflammatory chemokines in the host muscle decreases inflammatory players like fibro-adipogenic progenitors or Macrophages (M1 or M2) making a more permissive niche in the Bindarit treated host to allow for better engraftment and survival. It is also plausible that inhibition of inflammatory cytokines during the first week after injury has direct effects on the Hu-MuSC. For example, it is possible that inhibition of inflammatory chemokines reduces proliferation and migration of satellite cells, favoring differentiation or quiescence. This could be tested in future studies involving a greater area of muscle injury. Our observations are consistent with a recent study showing that inhibition of inflammatory-induced activation of CCL2 and CCR2 signaling after injury enhances regeneration and functional recovery^33^. Our findings suggest that at least part of the mechanism underlying that observation involves satellite cell mediated muscle formation.

There is evidence of chronic activation of aged satellite cells in mice^34^, suggesting senescence cellular programs involving inflammatory cytokines^35^. It is also known that a chronic inflammatory milieu (high IL-6, tumor necrosis factor-α (TNF-α), interleukin-1 (IL-1), and C-reactive protein (CRP) is correlated with sarcopenia in aging^36^. These cytokine profiles are thought to lead to a predisposition to age-related sarcopenia, ultimately leading to exhaustion of the PAX7+ satellite cells niche^36,37^. Our data shows that most of the inflammatory cytokines including IL-6, IL-33, IL-8, TNFRSF12A and chemokines CCL2, CCL7, CCL8, were upregulated in stimulated Hu-MuSC during activation. This suggests that similar gene expression changes that occur with acute injury and activation occur perhaps in a deregulated fashion in human muscle diseases and aging.

## Materials and Methods

### Isolation of Hu-MuSC

We have previously developed methods for isolation and purification functional satellite cells from human muscle tissue^11,13^. This preparation of purified and minimally altered satellite cells enables in vitro and in vivo experimentation as well as translational applications. Immediately after surgical resection the muscle is stored in Collection media composed of 30% FBS in DMEM high glucose, supplemented with Pen Strep at 4^Ο^C. The protocol for Human satellite cell activation begins 12 hrs before the start of processing the muscle biopsy, take 2/3 of the muscle tissue specimen and keep at 4^Ο^C for the quiescent sample. The remaining 1/3 of the muscle biopsy will be for the experimentally stimulated sample, and it will be immediately macrodissociated: a mechanical dissociation of the muscle sample is done using a blade. To ensure a clear single cell suspension we use enzymatic dissociation and filters as well as Magnetic Cell Separation (MACS LD) of hematopoietic cells (CD45+ and CD31+) from the cell suspension by means of antibodies and magnetic columns. To specifically sort MuSC, we used all of the following antibodies as muscle stem cells surface markers: CD56, CD29 and CXCR4. Please refer to our detailed published protocol for all details^12,13^. After this, resuspend the macrodissociated tissue in 30 ml of Growth Media (Ham’s F10 basal media, 20% FBS, 1X P/S, 1/500 1M HEPES, 5ng/ml FGF2) and transfer to a low attachment plate. Incubate at 37^Ο^C during 12 hrs. After this incubation, collect the macrodissociated muscle tissue from the plate and transfer to a 50 ml conical tube. Rinse plate with 1x PBS. Bring volume to 50ml with 1x PBS. Centrifuge at 2000 rpm for 5min. Aspirate supernatant and add digestion media and continue with the normal digestion/processing/staining protocol together with the quiescent muscle sample from the same individual^13^.

### Single cell Assay for Transposase Accessible Chromatin (scATAC) in Human Satellite Cells

All protocols to generate scATAC-seq data using the 10x Chromium platform, including sample preparation, library preparation and instrument and sequencing settings, are described below and are also available here: https://support.10xgenomics.com/single-cell-atac.

To investigate the differentially open chromatin regions in controls quiescent vs. activated Hu-MuSC after the overnight activation model, right after the FACS isolation, the nuclei were isolated following guidelines for limited number and fragile cells by 10x Demonstrated protocol (Nuclei Isolation for Single Cell ATAC Sequencing CG00169 Rev E) and the nuclei sample from quiescent and activated Hu-MuSC were loaded into the chromium chip by the UCSF Genomic core for scATAC.

The scATAC-seq libraries were prepared and sequenced according to the Chromium Single Cell ATAC Reagent Kits User Guide (10x Genomics; CG000168 Rev B) and the BCL files subjected to the Cell Ranger pipeline. (Cell Ranger Software v.1.2.0; https://support.10xgenomics.com/single-cell-atac/software/pipelines/latest/algorithms/overview).

### Data processing using Cell Ranger ATAC software 1.2.0

(https://support.10xgenomics.com/single-cell-atac/software/pipelines/latest/what-is-cell-ranger-atac) and Loupe Cell Browser Version 5.0.1 (https://support.10xgenomics.com/single-cell-atac/software/visualization/latest/what-is-loupe-cell-browser) were used as described before^38^. Analysis was done using R software^39^, the Seurat^40^ v4.3.0 and the Signac v1.9.0 packages^41^, designed as a framework for the analysis of single-cell chromatin data and related vignettes^41^. Single-cell chromatin state analysis with Signac^41^ was employed to enable end-to-end analysis of chromatin data and includes functionality for diverse analysis tasks, including identifying cells from background noncell-containing barcodes, calling peaks, quantifying counts in genomic regions, quality control filtering of cells, dimension reduction, clustering, integration with single-cell gene expression data, interactive genome browser-style data visualization, finding differentially accessible peaks, finding enriched DNA sequence motifs, transcription factor foot printing and linking peaks to potential regulatory target genes^42^. Next, we processed the gene expression assay by normalizing RNA counts with SCTransform (p<0.05) and Seurat. Significant peaks (p<0.05) allowed for gene enrichment identification and pathway analysis using the Reactome PA^43^.

### Data and materials availability

Single cell open chromatin fastq files and filtered matrices as well as detailed scripts, can be found here: https://github.com/kstriedinger/PomerantzLab_ATAC1.git

The open-source data repository that matches the single cell RNA dataset is https://datadryad.org/stash/dataset/doi:10.7272/Q65X273X. The code for the cytokines is like the source code described before^12^, using in this case the featured plots for the cytokines’ genes.

### Cytokine autoantibody panels

Hu-MuSC from one Vastus, one Pectoralis and one Rectus from humans ages 29, 31 and 44 years old were cryopreserved and were later thawed and plated on Matrigel precoated dishes in growth media (20% FBS in DMEM high glucose + P/S). After a media change 12 hrs later, Hu-MuSC were cultured under regular conditions for 3 to 6 days. Then the supernatant and the Hu-MuSC were harvested in separate samples and sent to Eve Technologies according to their protocol for sample collection. Undiluted supernatant and cell lysate samples were profiled for 83 different human cytokines using the Human Cytokine Array/Chemokine Array 71-Plex Panel and the 12-plex featured assay (catalog no: HD71; Eve Technologies). The controls included growth media and lysis buffer processed under the same conditions as the samples but without cells. Samples were prepared according to Eve Technologies® guidelines (https://www.evetechnologies.com/sample-collection-dilution-and-storage/) for the MILLIPLEX® Human Cytokine Autoantibody Panels and the magnetic microsphere beads from Luminex® Corp. Briefly, each set of beads is distinguished by different ratios of two internal dyes yielding a unique fluorescent signature to each bead set. Capture antibodies or antigens were coupled to the magnetic beads. Eve Technology results were based on the fluorescence intensity values in direct proportion to the standard known concentrations to calculate the concentration of cytokine proteins in our samples. Fluorescence values of the control supernatant or the control lysis buffer were subtracted from the fluorescence values of the supernatant sample or Hu-MuSC samples respectively.

### RT-QPCR

Total RNA was isolated using the RNAeasy isolation kit (Qiagen Cat # 74104). RNA was transcribed into cDNA with High-Capacity cDNA Reverse Transcription kit (ThermoFisher Scientific, Cat # 4368814). cDNA was then pre-amplified with GE PreAmp Master Mix (Fluidigm Inc, Cat#100-5876C2). Real-time quantitative PCR was performed in triplicated with either Taqman Universal PCR Master Mix (Life Technologies) or SybrGreen primers on a Viia7 thermocycler (Life Technologies). Taqman primers are listed in the Table 1 below. B-Actin, RPS-13 and GAPDH were used for normalization as endogenous control. Cytokine primers included:

**Table 1.**
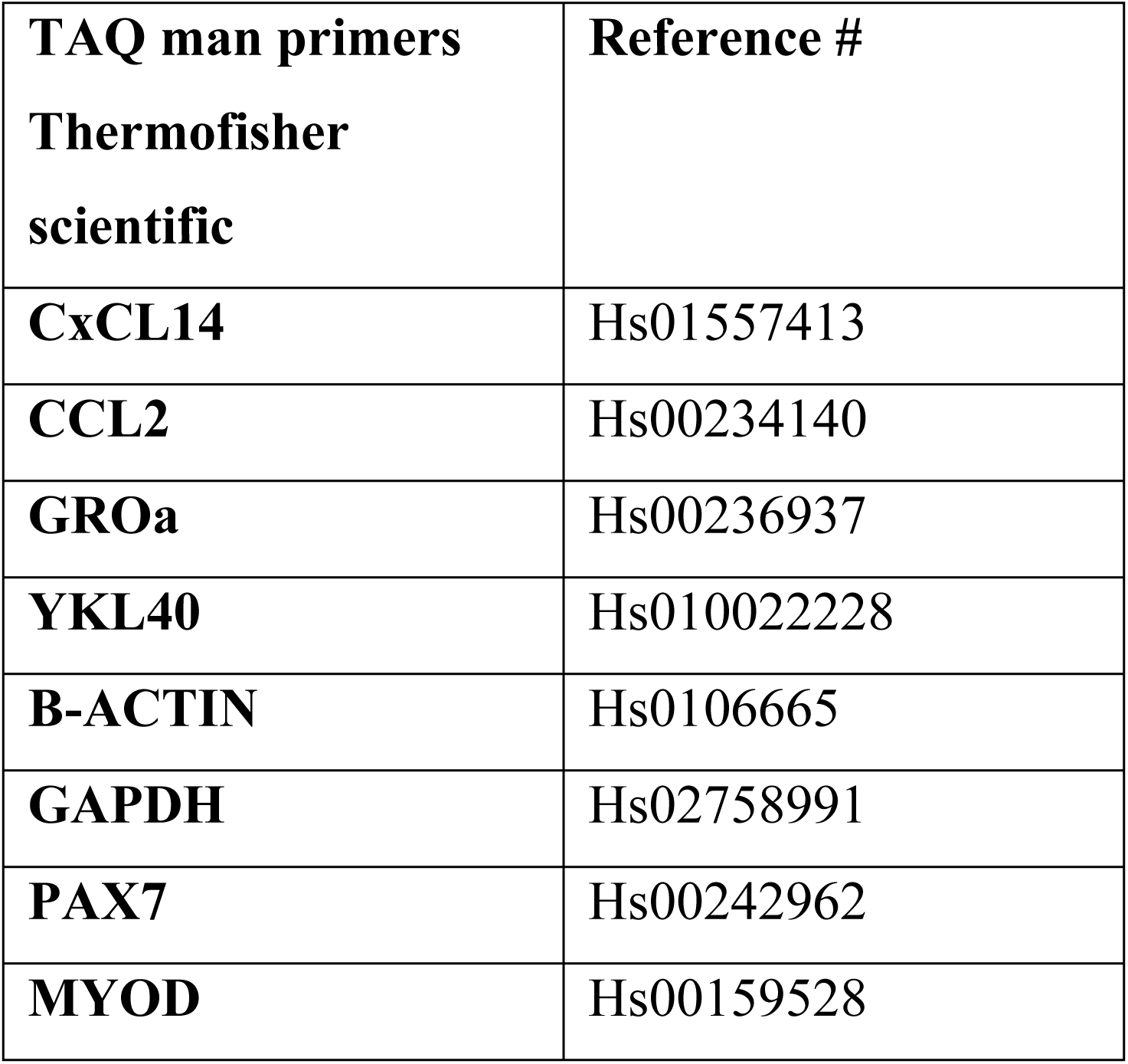
List of primers used in quantitative RT-PCR.

| **TAQ man primers**<br>**ThermoFisher**<br>**scientific** | **Reference #** |
| --- | --- |
| **CxCL14** | Hs01557413 |
| **CCL2** | Hs00234140 |
| **GROa** | Hs00236937 |
| **YKL40** | Hs010022228 |
| **B-ACTIN** | Hs0106665 |
| **GAPDH** | Hs02758991 |
| **PAX7** | Hs00242962 |
| **MYOD** | Hs00159528 |

To analyze RT-PCR data, Wilcoxon tests in Prism software were performed on 3 to 5 paired control and experimentally stimulated Hu-MuSC, each pair coming from one human donor muscle either vasti or rectus, p<0.05 as significant.

### Xenotransplants

2,000 Hu-MuSC were co-injected with Bupivacaine in the left TA of irradiated NSG mice (6-10 weeks old). To test the effects of cytokine’s on engraftment and myofiber formation of Hu-MuSC, we treated with a daily dose during 7 days of either, Bindarit - the selective inhibitor of CCL2, CCL7 and CCL8, or the Recombinant human CCL-2 from Preprotech 100 mg/kg, or vehicle only controls. Three weeks after Hu-MuSC transplantation, animals were sacrificed, and the TA was processed for Cryosections and immunostaining. Controls were the same vehicle, volume, administration route (IP) and frequency as the Bindarit group but without the Bindarit agent. Bindarit was chosen because it causes dose dependent selective inhibition against monocyte chemotactic proteins CCL2, CCL7, CCL8. Anti-inflammatory efficacy in animal disease models *in vivo* (100 mg/kg ip, rats & mice). Bindarit modulates cancer-cell proliferation and migration, mainly through negative regulation of TGF-β, AKT signaling and the NF-κB pathway. In our experiment we used Bindarit at 100 mg/kg IP BID. Three weeks after transplantation the TAs were collected in OCT, cryopreserved and cryosected. The number and diameter of the human fibers Dystrophin positive in the mouse muscle niche, as well as the Hu-MuSC expressing PAX7 positive Human Spectrin positive immunostaining were quantified.

Using three independent donors from which human muscle satellite cells were isolated in independent xenotransplant experiments, 2,000 Hu-MuSC were co-injected with Bupivacaine in the left TA of NSG mice (6-10 weeks old). Three weeks after Hu-MuSC transplantation, animals were euthanized, and the TA was processed for cryosections and immunostaining. We observe a higher number of human fibers identified by human Dystrophin immunostaining in the Bindarit treated group compared to controls. We also observe that the fibers in the Bindarit group have larger size compared to controls. These results support the idea that lowering the levels of cytokines in the host muscle niche, by means of the Bindarit, results in higher engraftment and myofiber formation capacity.

### Analysis of images in Fiji

Differential fluorescence labeling and multi-fluorescence imaging followed by Object-based colocalization analysis (OBCA) tools^44^.

### Ethics

#### Human subjects

This study was conducted under the approval of the Institutional Review Board at The University of California San Francisco (UCSF). Written informed consent was obtained from all subjects.

#### Animal experimentation

All procedures were approved and performed in accordance with the UCSF Institutional Animal Care and Use Committee (Protocols #181101).

